# Machine learning approach to predict protein-protein interactions between human and hepatitis E virus: revealing links to hepatocellular carcinoma

**DOI:** 10.1101/2025.02.23.639757

**Authors:** Anahid Hematpour, Parnian Habibi, Sajad Alavimanesh, Katayoon Dadkhah, Kiarash Babaie

## Abstract

Hepatitis E virus (HEV) remains a widespread yet underrecognized cause of acute and chronic liver disease, contributing to an estimated 44,000 to 70,000 deaths annually. With recurrent HEV outbreaks and limited treatment options, there is an urgent need for effective therapeutic development. Though, many aspects of the HEV life cycle, particularly the host-virus interactions that shape infection outcomes, remain poorly understood. Understanding virus-host protein-protein interactions (PPIs) is essential for targeted drug discovery, a task increasingly facilitated by advancements in machine learning. Here, we applied KNN, SVM, NV, and RF to predict novel HEV-human PPIs, offering critical insights into pathogen virulence strategies and potential therapeutic targets. Among 88 descriptors, the most effective features were GC content, Gene Ontology semantic similarity, Normalized frequency of beta-structure, and normalized frequency of alpha-helix and coil. Among the models, DT achieved the highest sensitivity (77%). while Logistic Regression (LR) had the highest specificity (52%) and the best accuracy of 0.61, showing robust prediction of positive and negative cases. Additionally, our proposed LR model has predicted novel potential targets in hepatitis E virus-human PPIs, which have been further validated through Gene Ontology enrichment analysis. Gene Ontology and disease enrichment analyses revealed HEV’s impact on immune modulation, lipid metabolism (FASN, APOB, EPHX2), and oncogenic pathways (FN1, JUN, HRAS, TP53), supporting its potential role in liver pathology and hepatocellular carcinoma (HCC). These findings provide novel insights into HEV-host interactions, offering targets for future antiviral strategies.

## Introduction

Hepatitis E virus (HEV) is a leading cause of acute viral hepatitis, responsible for 3.3% of viral hepatitis-related deaths with the estimated fatalities ranged from 44,000 to 70,000 (1, 2). Globally, HEV infection is on the rise (3–5), with around 20 million new cases annually, 20% of which are symptomatic (6, 7). In a Scottish study involving 3,500 individuals with suspected acute viral hepatitis, HEV detection rates were 31 times higher than Hepatitis A and seven times higher than Hepatitis B, highlighting its increasing significance as a cause of acute hepatitis (4). These estimates rely on studies using assays now recognized for their limited sensitivity, suggesting that the actual global burden of hepatitis E-related disease is likely underestimated (8).

HEV is a quasi-enveloped, icosahedral, positive-sense, single-stranded RNA virus in the Paslahepevirus genus of the Hepeviridae family (1). Among the eight genotypes in Orthohepevirus A, GT1–4 are most relevant to humans. HEV primarily targets hepatocytes but also affects the small intestine, colon, and neuronal system, linking it to neurological (e.g., Guillain-Barré syndrome, encephalitis) and renal complications in up to 16% of symptomatic patients (1, 2, 8). HEV infections exhibit marked global variability in distribution and transmission. In low-income countries, genotypes 1 and 2 are transmitted via contaminated water, causing outbreaks with mortality rates of 0.2–4 Genotypes 3 and 4, prevalent in high-income nations, are zoonotic, linked to undercooked meat, and typically asymptomatic; however, symptomatic cases can involve severe illness, with jaundice (40%), hospitalization (75%), and mortality (3%), especially in those with chronic liver disease (1, 7). Medical interventions such as blood transfusion and organ transplantation also contribute to HEV transmission (9).

Global seroprevalence varies widely from 27–80% in endemic areas (e.g., India, Southeast Asia) to 2–20% in non-endemic regions (e.g., Europe, the USA, East Asia). A meta-analysis estimated a global prevalence of 12.47%, with the highest rates in Africa (21.76%) and Asia (15.80%) (7). Individuals with chronic liver disease, diabetes, obesity, immunosuppression, and other comorbidities face heightened susceptibility to severe HEV infections, with increased mortality (2, 7, 9). Pregnant women, particularly in the third trimester, are at significant risk of acute liver failure and pregnancy complications, including fetal loss and maternal mortality rates of 5.1–31% (2, 7, 10). In 2005, HEV was linked to over 70,000 deaths and 3,000 stillbirths worldwide (2). Despite its global burden, HEV remains under-prioritized in public health. Of 44 documented outbreaks across 19 countries, 20% are unreported in peer-reviewed literature. Most occurred in Africa (61.4%) and Southeast Asia (27.3%), mainly in humanitarian settings, hospitals, and workplaces. Genotype 4 outbreaks (9.1%) were limited to high-income countries, linked to contaminated food (5). The periodic nature of outbreaks, driven by declining anti-HEV IgG seroprevalence, highlights the urgent need for new therapies (8).

There is evidence suggesting that HEV may accelerate the progression of chronic liver disease to hepatocellular carcinoma (HCC), often associated with cirrhosis. A large US study found HEV IgG infections associated with increased fibrosis risk, while an Eastern Chinese study showed higher anti-HEV antibody prevalence in cancer patients, especially those with leukemia (32.3%) and liver cancer (31.1%), compared to controls (13%). A systematic review and meta-analysis by Yin et al. indicated that HEV infection (11) increases the risk of HCC.

There is no approved specific antiviral for HEV, and vaccination is available only in China for genotype 1. Current treatments for hepatitis E, including ribavirin and PEGylated interferon-α (PegIFN-α), are limited and pose challenges. Ribavirin, though relatively effective, is contraindicated during pregnancy due to teratogenicity, and some patients do not respond or remain viremic despite long-term use. Additionally, ribavirin-associated mutations may increase HEV replication, worsening outcomes. PegIFN-α, due to its immunostimulatory properties, cannot be used in transplant patients, especially those with heart, lung, pancreas, or kidney transplants, as it risks acute rejection. (7, 12).

Understanding infection mechanisms, host immune responses, and efficient molecular targeting recognition for the development of therapeutics relies heavily on identification of functional protein-protein interactions (PPIs) (13). PPIs, representing the initial contact between viral proteins and host receptors, are a key focus of research (10,11). Advances in computational methods, including machine learning algorithms, have made predicting and validating virus-host PPIs more feasible in recent decades, surpassing time- and labor-intensive wet lab methods, which are prone to false-negative results. Specifically, machine learning techniques have demonstrated exceptional ability in recognizing patterns for sequence-based biological prediction tasks (14–19).

Despite extensive research on viral hepatitis, human-HEV protein-protein interactions remain largely uncharted. Barman et al. (2014) developed a supervised machine learning-based approach to predict HBV/HEV-human PPIs, integrating domain-domain association, network topology, and sequence features. While their study provided a framework for predicting unknown HEV-human PPIs, the evaluation of HEV-specific interactions remained limited, and performance metrics for HEV-host interactions were not explicitly reported. To build upon these findings, we present a refined approach that specifically focuses on HEV-host interactions, utilizing classifiers including Logistic Regression, SVM, Naïve Bayes, Decision Trees, Random Forest, and KNN using metrics like sensitivity, specificity, accuracy, PPV, NPV, AUC, and prevalence rate. Key predictors such as Amino Acid Indices in Aaindex1, Amino acid sequence-based features, Nucleotide sequence-based features (20), Gene Ontology semantic similarity, and Network topology-based features. This study refines HEV-host PPI prediction, addressing previous gaps and advancing HEV pathogenesis understanding and therapeutic target identification.

## Material and methods

### Dataset

To perform a virus-host PPI classification task effectively, both positive (existing interactions) and negative (non-interactions) samples need to be learned or considered for training a predictive model.

### Positive dataset

To generate positive host-viral protein-protein interactions (HI-PPIs) (21), all interactions involving HEV were collected from databases: Intact, Virus Mint, DIP, STRING, and BioGRID (22).

### Negative dataset

Creating a reliable negative dataset for PPI prediction is challenging due to the lack of experimentally verified non-interacting protein pairs. Previous studies suggest that interspecies PPI prediction requires a balanced ratio of positive to negative samples (1:1), but standardized negative samples are lacking. Drawing on viral-host protein pathway research, which shows viruses target highly connected human proteins, we designed the negative dataset using the Human Protein Reference Database (HPRD) release 9, following Chakraborty et al. After excluding proteins common to the positive dataset, we identified unique, non-redundant human proteins, ranked by interaction degree. Proteins with lower connectivity were selected as negative dataset candidates, and sequences for the proteins with the lowest interaction degrees were extracted from Uniprot database (23). Positive and negative datasets were then divided into training and independent sets. Specifically, 80% of total protein pairs from each dataset were randomly selected for training, while the remaining pairs formed the independent datasets.

### Encoding proteins as feature vectors

The performance of classification algorithms depends on effective feature extraction. Since proteins vary in length, consistent representation of protein sequences is crucial for applying machine learning to PPIs (17, 23).

To represent human and HEV proteins as feature vectors, we utilized the AAindex database, which provides numerical indices for various physicochemical and biochemical properties of amino acids. The AAindex is divided into three sections: AAindex1 (amino acid indices with 20 numerical values), AAindex2 (amino acid substitution matrix), and AAindex3 (statistical protein contact potentials). Key physicochemical properties include hydrophobicity, alpha propensity, and beta propensity. The AAindex comprises 544 sets of physicochemical properties from various literature sources, with a selected subset used as features for the classification task (24).

### Amino Acid Indices in Aaindex1

#### Physicochemical indices

- **STERIMOL length of the side chain**: STERIMOL constants contribute to exploring structure-activity relationships in peptides (25) and describe the steric bulk of a substituent based on its dimensions in three spatial directions. “L” denotes the length of the side chain, measured in the direction in which it is attached to the glycine backbone (26, 27).
- **Volume:** Molecular volume stands out as one of the properties most strongly correlated with protein residue substitution frequencies. (28).
- **Van der Waals parameter R_0_:** The empirical measure of the atomic size, representing the distance between the centers of adjacent atoms when they are just in contact. It varies for different atoms within the amino acid, reflecting their specific van der Waals radii (29).
- **Alpha-CH chemical shifts:** These chemical shifts offer insights into the local structural environment, aid in determining secondary structure elements, and help map interaction sites. Changes in chemical shifts can indicate conformational changes upon interaction, helping to understand the dynamic behavior of proteins during interactions (30).
- **Mean volumes of residues buried in protein interiors:** It involves measuring and analyzing the average volume occupied by amino acid residues that are located within the interior of a protein, providing insights into the packing and compactness of the protein’s core and shedding light on the factors influencing protein folding, stability, and structure. The mean volumes of buried residues are calculated based on the three-dimensional coordinates of protein structures and contribute to our understanding of the structural features crucial for maintaining a protein’s three-dimensional architecture (31).

#### Hydrophobicity indices

- **The hydrophobic component of transfer free energies for amino acid side chains in α-helical polypeptides:** It presents the values for the hydrophobic component of the free energy of water-oil transfer for different amino acid side chains in an alpha-helical conformation. It is calculated based on the surface area and the hydrophobicity of each amino acid. This calculation takes into account the idea that hydrophobic amino acids prefer a nonpolar environment (like oil) over water (32).
- **Free energy in beta-strand conformation:** It refers to the energy associated with the stability of amino acids adopting a beta-strand structure in proteins. It quantifies the thermodynamic preference of amino acids for forming beta-strands, as indicated by their positions in the Ramachandran plot not considering the identity and the conformation of the surrounding residues in the amino acid sequence (33).
- **Scaled side chain hydrophobicity values:** The hydrophobic characteristics of the side chains of the 20 common physiological amino acids in proteins that undergo post-translational or co-translational modifications (34).
- **Optimized relative partition energies - method D:** It is a computational approach for predicting PPIs and assessing complex stability. It calculates Relative Solvent Accessibility (RSA) values for amino acids, introduces optimized parameters for each amino acid type, and computes ORPE to represent individual amino acid contributions to complex stability based on solvent exposure. The sum of ORPE values predicts interactions, where lower total energies indicate more stable interactions. Method D refines earlier methods (A-C) with optimized parameters, enhancing accuracy in predicting protein-protein interaction energies and complex stability (35).
- **Hydration potential:** Refers to the energetic considerations associated with the transfer of a nonpolar solute from a hydrophobic environment to an aqueous (polar) environment. This concept is crucial for understanding the hydrophobic effect, a phenomenon in which hydrophobic molecules or residues tend to cluster together to minimize their exposure to water, contributing to the stability of biomolecular structures(36).
- **Mean fractional area loss (MFAL):** MFAL is a parameter used to quantify the packing efficiency of amino acid side chains within the core of a protein. It assesses how much space is lost or excluded in the protein interior due to the presence of side chains. A lower MFAL indicates a more tightly packed hydrophobic core, which is generally associated with a well-folded and stable protein structure (37).
- **Flexibility parameter for one rigid neighbor:** Component of the Karplus–Schulz matrix used in predicting torsion angles (phi, psi, and omega) in proteins. It represents the influence of the dihedral angle of one residue on its neighbor in the presence of one neighboring residue. The Karplus–Schulz model is a statistical approach that correlates dihedral angles with local protein structure, and this flexibility parameter captures the energetic and geometric effects of neighboring residues on torsional angles (38).
- **Normalized flexibility parameters (B-values), average:** Also derived from the Karplus and Schulz approach, it represents a measure of the flexibility or mobility of amino acid residues within protein structures. These B-values are normalized to have a mean of 1.0 and a root mean square deviation of 0.3, facilitating consistent comparison across different proteins. Residue types are categorized as flexible or rigid based on their average normalized B-values, with values below 1.0 indicating rigidity (39).
- **Ratio of buried and accessible molar fractions:** It is a metric used to quantify the distribution of amino acid residues in proteins. It compares the molar fractions of a specific amino acid type that is either buried within the protein’s interior or accessible on its surface to the solvent. The ratio provides insights into the preference of a particular amino acid type to be buried or exposed, offering valuable information about the protein’s structural characteristics, hydrophobicity, and packing (40).
- **Atom-based hydrophobic moment:** Quantifies the distribution and orientation of hydrophobic residues within a protein structure at the atomic level by considering their individual contributions and spatial arrangement. This concept provides insights into the anisotropy of hydrophobicity in proteins and has been used to study aspects of protein folding, stability, and structure prediction (41).
- **Residue accessible surface area (ASA) in folded protein:** ASA in folded proteins refers to the surface area of an amino acid residue that is accessible to solvent molecules in the folded state of a protein. It is calculated by considering the difference between the total surface area of the residue and the surface area buried upon protein folding. Residue ASA is crucial for understanding the solvent exposure and packing of amino acid residues within a protein, providing insights into protein structure, function, and interactions (42).
- **TOTFT (Total Polarity Index of Face Turns) and TOTLS (Total Polarity Index of Loop Structures):** These indices were designed to capture various aspects of protein structure and behavior, with a focus on hydrophobicity, polarity, and structural features like turns and loops. TOTFT is the sum or combination of the primary and alternative polarity indices for face turns in a protein. TOTLS is the sum or combination of the primary and alternative polarity indices for loop structures in a protein (43).
- **Optimal Matching Hydrophobicity:** Prediction of transmembrane segments in membrane proteins. Membrane-spanning regions often consist of hydrophobic alpha helices, and this method aimed to identify such patterns within protein sequences (44).
- **Transfer Free Energy from Octanol to Water:** This term refers to the free energy change associated with transferring a solute molecule from an octanol (hydrophobic) environment to water (hydrophilic) environment. It quantifies the energetics of a solute’s interaction with hydrophobic and hydrophilic environments, providing insight into the hydrophobic effect, which plays a significant role in processes such as protein folding and ligand binding (45).
- **Weights for beta-sheet at the window position of -5:** Assigns different weights to amino acid positions within a window surrounding a particular residue in a protein sequence for predicting secondary structure elements, such as beta-sheets or coil regions. This weight reflects the importance of the amino acid residues at -5 positions in contributing to the prediction of beta-sheet secondary structure. The positive or negative weights would indicate the degree of influence each position has on the prediction.
- **Weights for coil at the window position of -6:** This is a weight assigned specifically to the amino acid position at -6 within a sequence window. It captures the contribution of the amino acid at position -6 in predicting coil regions in the secondary structure (46).
- **Transfer free energy to surface:** It refers to the energy required or released when amino acid residues move from a solution to the surface. This transfer free energy to the surface serves as a measure of the hydrophobicity of amino acids, with lower surface tension values indicating higher hydrophobicity (47).
- **Normalized positional residue frequency at helix termini N’:** It refers to a measure in the context of α-helices, indicating the preference of specific amino acid residues at the N-terminus (NH₂-exposed end) and C-terminus (COOH-exposed end) positions. It involves comparing the occurrence of a residue at N’ position to its overall frequency in the entire dataset, with values greater than 1 indicating a preference and values less than 1 indicating a dispreference. The term is used to quantify how certain residues are favored or avoided at the ends of helices, considering disruptions in the regular hydrogen bonding pattern (48).
- **Loss of Side chain hydropathy by helix formation:** It refers to the reduction in amino acid side-chain hydrophobicity when an extended polypeptide folds into an α-helix. It’s estimated by comparing hydrophobic components in extended and helical conformations, revealing an average loss of about 0.6 kcal/mol per residue. The resulting hydropathy scale is a rough approximation of the most favorable transfer free energies during helix formation, considering exceptions for specific amino acids and potential corrections for hydrogen bonding effects (49).
- **Free energies of transfer of AcWl-X-LL peptides from bilayer interface to water:** Energy changes during peptide transition from the lipid bilayer interface to water. These values contribute to an interfacial hydrophobicity scale, offering insights into peptide-membrane interactions. Despite approximations, the scale serves as a crucial reference for understanding the thermodynamics of these interactions (50).

#### α-propensity indices

- **Normalized frequency of middle helix:** It is a measure that standardizes the occurrence of amino acids in the middle residues of helices across different proteins, correcting for variations in amino acid composition (51).
- **Helix initiation parameter at posision i,i+1,i+2**: It refers to a parameter used for describing the initiation of α-helical structures in peptides. It considers specific interactions involving side chains, the N- and C-termini of helices, and influences the statistical weight of an α-helix between specified positions in a peptide. This parameter contributes to determining the likelihood of α-helix formation at the designated positions within the peptide sequence (52).
- **Helix formation parameters (delta delta G):** It measures the energy change when a peptide transitions from a random coil to a helical structure. It indicates the thermodynamic stability of helices, comparing the free energy of helix formation for a specific sequence to a reference state like glycine (Gly), revealing the intrinsic conformational preferences (53).
- **Thermodynamic beta sheet propensity:** It measures how amino acids preferentially contribute to the stability of antiparallel β-sheet structures, determined by their substitutions in a zinc-finger host peptide. This scale is established through precise energy measurements using a competitive cobalt(II)-binding assay in aqueous solution (54).
- **Alpha helix propensity of position 44 in T4 lysozyme:** It indicates the likelihood of amino acids, when substituted at this site, to influence the formation of an alpha helix. The measurement involves assessing stability changes resulting from amino acid substitutions at the solvent-exposed position within the alpha helix (residues 39 to 50) in T4 lysozyme. This provides insights into the structural preferences of various amino acids at position 44 regarding their impact on alpha helix formation (55).
- **Normalized positional residue frequency at helix termini N3 and C’ (Aurora-Rose):** It shows the frequency of a particular amino acid at the N-terminus or C-terminus of an alpha-helix, specifically at the N3 position, normalized against its overall distribution in the dataset. A normalized frequency of 1 implies no preference, while values above or below 1 signify selection for or against that residue at the C-terminus or N3 position of the N-terminal helix (48).
- **Hydrophobicity Coefficient (RP-HPLC, C4, 0.1%TFA/MeCN/H2O):** A measure of amino acid or peptide hydrophobic affinity in C4 reversed-phase high-performance liquid chromatography (RP-HPLC), using a mobile phase of 0.1% trifluoroacetic acid, acetonitrile, and water. This coefficient aids in characterizing the hydrophobic contribution to protein stability, offering insights into surface free energy, protein folding, and interactions that influence protein-protein binding (56).
- **Linker propensity from medium dataset (linker length is 6--14):** Linker propensity in the medium dataset refers to the preference of specific amino acids within linkers of moderate length (6-14 residues). It involves calculating the ratio of amino acid occurrences in the linker set to their occurrences in the full protein set (57).
- **Principal component IV:** It is a composite chemical factor derived from the physico-chemical properties of amino acids. It appears to involve hydroxyl and sulphydryl groups, along with a potential connection to the ability of amino acids to form hydrogen bonds. It contributes to understanding the relationship between chemical structure and biological activity in hormone peptides (58).
- **Normalized frequency of alpha-helix and coil:** The normalized frequency of alpha-helix is the percentage of amino acid residues predicted or observed to form alpha-helical structures in a protein sequence. Similarly, the normalized frequency of coil represents the proportion of residues in coil conformations. Both frequencies are expressed relative to the total sequence length (59).
- **Information measure for coil, loop, and turn:** Information measure for coil, loop, and turn refers to a quantitative assessment of the relationship between amino acid sequence and the conformational states of coil, loop, and turn structures in globular proteins. These metrics provide a means to systematically assess how certain amino acid sequences contribute to the structural characteristics of coil, loop, and turn regions (60).

#### β-propensity indices

- **Normalized frequency of beta-structure:** It denotes a scaled measure quantifying the prevalence of beta-structures across diverse proteins. The term implies an adjustment for comparability (59).
- **Information measure for pleated-sheet:** It quantifies the significance of residues adopting pleated-sheet conformation in proteins. It assesses the frequency of pleated-sheet occurrences at various residue separations (m values), representing the distances between amino acid residue pairs. This measure offers insights into amino acid preferences within these structures and contributes to statistical analyses of protein sequences and conformations, considering different separation distances along the protein chain (60).
- **Conformational preference for all beta-strands:** It indicates the likelihood of amino acid residues adopting specific structural arrangements within antiparallel β-sheets, with a focus on hydrogen bond patterns and chain connectivity (61).
- **Propensity of amino acids within pi-helices:** It refers to the specific amino acid preferences within π-helical structures. Aromatic and large aliphatic amino acids are favored, while small amino acids like Ala, Gly, and Pro are avoided. These propensities are distinct to π-helices and significantly differ from overall amino acid distributions (62).

#### Composition indices

- **AA composition of CYT2 of single-spanning proteins:** Distribution of amino acids within the cytoplasmic region (CYT2) of type-11 single-spanning membrane proteins. Analyzing the types and proportions of amino acids in this region provides insights into the chemical makeup of the cytoplasmic side in single-spanning membrane proteins.
- **AA composition of EXT of Multi-spanning proteins:** It describes the amino acid distribution in the extracellular region of multi-spanning proteins, revealing the composition of the portion facing the external environment.
- **AA composition of CYT of multi-spanning proteins:** It refers to the amino acid distribution in the cytoplasmic region of multi-spanning proteins, providing insights into the chemical composition on the cytoplasmic side.
- **AA composition of MEM of single-spanning proteins:** It details the amino acid distribution in the transmembrane region of single-spanning proteins, indicating the composition of the portion crossing the cell membrane (63).
- **Distribution of amino acid residues in the 18 non-redundant families of mesophilic proteins:** It refers to the relative occurrence or arrangement of individual amino acids within the protein sequences of 18 distinct families of mesophilic proteins. This analysis is crucial for understanding the unique amino acid patterns in proteins adapted to moderate temperatures, offering insights into factors influencing protein stability and function. Comparing these distributions with thermophilic proteins can reveal molecular mechanisms relevant to stability (64).
- **Entire chain composition of amino acids in extracellular proteins of mesophilic bacteria:** Overall distribution and abundance of amino acids throughout the entire amino acid sequence of extracellular proteins in mesophilic bacteria. This comparison provides insights into how proteins have adapted to different environmental conditions, revealing selective pressures, functional adaptations, and structural changes (65)

Other features applied in this study are discussed below:

#### Amino acid sequence-based features

- **Amino acid composition (ACC):** Amino acid composition is the percentage of each amino acid in a protein.
- **Conjoint Triad (CT):** The conjoint triad (CT) feature groups three adjacent amino acids into units and calculates their frequency in a protein sequence. CT categorizes amino acids into seven groups based on their properties. Using a 3-amino acid window, the frequency of each CT is determined as the window slides through the sequence from N- to C-terminus, resulting in a 686-dimensional vector representing a protein pair.
- **Relative frequency of amino acid triplets (RFAT):** It quantifies the prevalence of specific amino acid triplets in proteins involved in PPIs. RFAT can provide insights into the functional relevance of certain amino acid combinations in the context of these interactions. It also allows us to assess the overrepresentation or underrepresentation of specific amino acid triplets in the proteins involved in host-pathogen interactions. Mathematically, RFAT is calculated as follows: RFAT(i, j, k) = (Count of the triplet “ijk” in the dataset) / (Total number of triplets in the dataset). “i,” “j,” and “k” represent the individual amino acids within the triplet.
- **Frequency difference of amino acid triplets (FDAT):** FDAT is a metric used to discover unique amino acid triplet patterns associated with PPIs. The formula for FDAT is as follows: FDAT (i, j, k) = RFAT(i, j, k) in interacting proteins- RFAT(i, j, k) in non-interacting proteins. “i,” “j,” and “k” represent individual amino acids within the triplet. By subtracting these relative frequencies, FDAT helps identify amino acid triplet patterns that are more or less common in host-pathogen interactions. This aids in pinpointing specific triplet sequences with potential functional relevance in the context of these interactions (66).
- **Pseudo amino acid composition (PAC):** PAC is utilized for predicting protein subcellular localization and membrane protein type. In contrast to amino acid composition, PAC relies on sequence-order information. It captures the frequency of individual amino acid composition while incorporating sequence-based data into its pseudo components (22).
- **Biosynthesis energy:** Amino acids are synthesized from metabolic precursors, such as pyruvate and 3-phosphoglycerate. The total cost of this process, referred to as biosynthesis energy, is calculated using the Wagner method (22).
- **Thermophilic propensity:** It refers to scores assigned to each amino acid, indicating their likelihood of contributing to the thermophilic nature of proteins. Calculated using the SCMTPP predictor from a dataset of both thermophilic proteins (TPPs) and non-TPPs, these scores are analyzed to better understand the physicochemical properties associated with TPPs (67).
- **Antegenic propensity:** Antigenic propensity is a calculated value indicating the likelihood of an amino acid being part of an antigenic determinant. It is derived from a method using physicochemical properties and epitope frequencies. Hydrophobic residues like Cys, Val, and Leu, when on the protein’s surface, tend to have higher antigenic propensities (68).
- **Disordered proteins:** Intrinsic disorder in proteins refers to regions that prevent them from adopting a specific three-dimensional structure but enable conformational changes during interactions. Pathogen-interacting proteins, especially those in host-pathogen networks, exhibit a higher proportion of intrinsically disordered regions. Proteins with intrinsically disordered regions are more prone to pathogen attacks (69).

#### Nucleotide sequence-based features

- **GC content:** The percentage of guanine and cytosine nitrogenous bases in a DNA molecule. This measure indicates the stability of DNA sequences because the guanine-cytosine bond forms a triple bond, making sequences with higher GC content more stable than those with lower GC content, where adenine and thymine form a double bond.
- **Codon usage:** Also known as codon bias, indicates how often different synonymous codons appear in coding DNA (70). It is determined by the ratio of the frequency of a specific codon (designated as “fi”) for a particular amino acid (indexed by “j”) to the total occurrence of that amino acid in the given sequence (referred to as “nj”). Mathematically, codon usage for the ith codon can be expressed as: *CU*i = fi / nj
- **Relative synonymous codon usage (RSCU):** Measure of codon usage that calculates the frequency of a specific codon in relation to the frequency that would be expected if all codons for the same amino acid were equally distributed. Mathematically, RSCU for the ith codon of the jth amino acid is computed as: RSCUi,j = (fi,j) / [(1/nj) * Σ(fi,j)] Here, fi,j represents the frequency of the ith codon for the jth amino acid, and nj is the total occurrence of that amino acid in the sequence and normalizes the codon frequency by dividing it by the expected frequency assuming equal distribution of codons.
**The Codon Adaptation Index (CAI):** A straightforward measure of the bias in Relative Synonymous Codon Usage (RSCU) and is calculated as follows: CAI = Π (RSCUi / RSCUmax) Here n is the length of the protein sequence, RSCUi is the RSCU value of the ith codon, and RSCUmax is the maximum RSCU value among codons for the amino acid associated with the ith codon. The CAI calculates the geometric mean of the RSCU values for each codon in a protein sequence and normalizes it by dividing by the geometric mean of the maximum RSCU values for each amino acid. This index is a useful indicator of how well the codon usage in a gene matches the codon usage in a reference set, often chosen to represent highly expressed and efficiently translated genes.
- **Stacking energy:** According to the nearest-neighbor (NN) model for nucleic acids, takes into account how the neighboring base pairs’ identity and orientation influence the stability of a particular base pair. The calculation of stacking energy (∇*G*) is as follows:

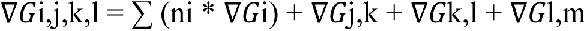

Here, ∇*G* for the initial (i), middle (j), and end (k, l) is determined using unified nearest-neighbor (NN) free energy parameters. If the duplex is self-complementary, the symmetry is maintained by setting ∇*G*l,m to +0.43 kcal/mol; otherwise, it’s set to zero if the duplex is non-self-complementary.
- **The interaction energy:** A measure of the dispersion and repulsion energies between a codon and its complement. It is calculated using the formula: IE = ∑ (ei * ni) / ∑ (ni) In this formula: IE represents the interaction energy, n is the length of the protein, ni is the frequency of the ith amino acid, and ei is the interaction energy associated with the ith amino acid. The interaction energy is determined by considering the frequency of each amino acid and its associated interaction energy.

### Gene Ontology semantic similarity

Gene Ontology (GO) provides a hierarchical framework to annotate genes across organisms, categorizing them by molecular function (MF), cellular components (CC), and biological processes (BP) (71). This structured approach aids in predicting protein-protein interactions (PPI), as interacting proteins often share similar biological roles, functions, and cellular locations, exhibiting high semantic similarity in GO terms (22).

#### Network topology-based features

- **Degree:** The degree of a protein refers to how many other proteins it interacts with.
- **Neighborhood Connectivity:** It is the average degree of all neighboring proteins of a given protein, excluding self-interactions.
- **Average Shortest Path Length:** This measures the average length of the shortest paths between the given protein and all other proteins.
- **Stress:** Stress represents the number of shortest paths passing through a specific protein in the human protein-protein interaction network (PPIN). It signifies the workload carried by that protein in the network.
- **Eccentricity:** Eccentricity measures the greatest distance between a specific protein and any other protein within the human PPIN (72).
- **Closeness:** Closeness is the reciprocal of the total distance from the given protein to all other proteins in the human PPIN.
- **Betweenness:** This centrality metric is a semi-normalized version of stress centrality. It calculates the ratio of the number of shortest paths passing through the protein to the total number of shortest paths between all protein pairs in the human PPIN.
- **Radiality:** Radiality is computed by subtracting the average shortest path length of a protein from the length of the longest path in the connected component, plus 1. To normalize it, the radiality of each protein is divided by the length of the longest path in the connected component.
- **Clustering Coefficient:** This coefficient measures the density of connections in the local region of a protein. It is the ratio of edges between a protein’s neighbors to the total possible edges among them (22).

### Post-translational modifications

Post-translational modifications (PTM) play a vital role in regulating protein functions, impacting PPIs in particular. To emphasize the significance of PTMs, we employed PTM such as deacetylation, phosphorylation, glycosylation, among others that are documented in the HPRD database to represent human proteins. For each set of 20 amino acids, the potential for 31 different PTM types exists. Consequently, each human protein was depicted as a binary feature vector with a length of 620. Each element within this vector, for a given protein, signifies whether a specific amino acid has undergone a specific PTM or not (28).

### Prediction algorithm

Common automated machine learning techniques (73–75), such as Random Forest, Naïve Bayes, and SVM, are often utilized to predict PPIs. These methods typically employ a five-fold cross-validation approach to evaluate their predictive performance. In this paper we took advantage of Linear regression (LR), K Nearest Neighbor (KNN), Support Vector Machine (SVM), Random Forest, and Naïve Bayes to build our models.

In brief, Support Vector Machine (SVM) is a powerful learning algorithm known for finding effective separation hyperplanes, making them suitable for linear classification. Random Forest is a popular supervised machine learning algorithm. It creates multiple decision trees and combines their outputs through majority voting for classification and averaging for regression, providing stability and accuracy in a wide range of machine learning applications. The Naïve Bayes algorithm, based on Bayes’ theory, is also a supervised learning approach primarily used for classification. It’s called “naïve” because it operates on the assumption that the value of a specific feature is independent of the value of any other feature (17, 22).

### Performance assessment measures

To evaluate feature selection’s impact, we used accuracy (AcC), sensitivity (Sen), specificity (Spec), positive predictive value (PPV), negative predictive value (NPV), and prevalence rate. TP and TN represent correctly predicted positive and negative cases, while FP and FN are misclassified instances. Accuracy reflects overall prediction correctness, sensitivity measures true positive identification, and specificity assesses true negative recognition (76). PPV indicates the proportion of true positives among predicted positives, while NPV measures true negatives among predicted negatives. Prevalence rate represents the proportion of actual positive cases in the dataset. These metrics collectively assess classification performance (16).

### Feature importance

To compute the contribution of the different descriptors in prediction, we removed each feature type in turn and then computed the accuracy of the proposed prediction model, the higher the loss of accuracy, the more important the feature.

### Disease Enrichment Analysis

To investigate whether the unknown virus-interacting proteins predicted in our study are associated with different stages or subtypes of liver cancer, the extracted gene set was submitted to Enrichr, selecting DisGeNET as the reference database. DisGeNET provides curated and predicted gene-disease associations, allowing us to determine whether the enriched proteins overlap with known liver cancer-associated genes (77).

Significantly enriched disease terms (p ≤ 0.005, adjusted for multiple testing) were examined for their biological relevance. If liver cancer-related terms were enriched, this would suggest that viral proteins interact with host factors involved in cancer-related pathways, potentially contributing to tumorigenesis or disease progression.

### GO annotation and pathway enrichment analysis

We further performed functional enrichment analysis to delineate genotype-specific host pathways targeted during infection, providing insights into viral adaptation and potential therapeutic targets. The DAVID web server (The Database for Annotation, Visualization and Integrated Discovery) was utilized to identify significantly enriched Gene Ontology (GO) annotation terms in the predicted human proteins interacting with hepatitis E virus. Gene Ontology (GO) term selection was performed across Molecular Function (MF), Biological Process (BP), and Cellular Component (CC) categories. We included only GO biological process annotation terms with a level greater than 2 and a significant false discovery rate (FDR) below 0.005.

## Results

Over all, we identified 196 interactions, all within the Hepeviridae family and specifically the Paslahepevirus genus. Notably, HEV genotype 1 (India/Hyderabad) exhibited the highest number of interactions (117), followed by HEV genotype 3 (Swine/US) with 68 interactions and HEV genotype 1 (China/HeBei) with 11 interactions.

### Classifier Evaluation Metrics Comparison

The bar plot (Figure 1) compares the performance of six classifiers (SVM, DT, KNN, RF, NB, and LR) across multiple evaluation metrics, including accuracy, sensitivity, specificity, positive predictive value (PPV), and negative predictive value (NPV). Among the models, DT achieved the highest sensitivity (77%). while LR had the highest specificity (52%) and the best accuracy of 0.61. Logistic Regression (LR) also led with an AUC of 0.63. PPV and NPV: Both were highest for Logistic Regression (LR), showing robust prediction of positive and negative cases. The other models, including Support Vector Machine (SVM), Naive Bayes (NB), and Random Forest (RF), showed moderate but less consistent performance, with accuracy values ranging from 0.56 to 0.60. Overall, while LR stands out as the most balanced model, DT demonstrated the highest sensitivity (0.77), making it more efficient in detecting true PPIs.

**Figure 1.**
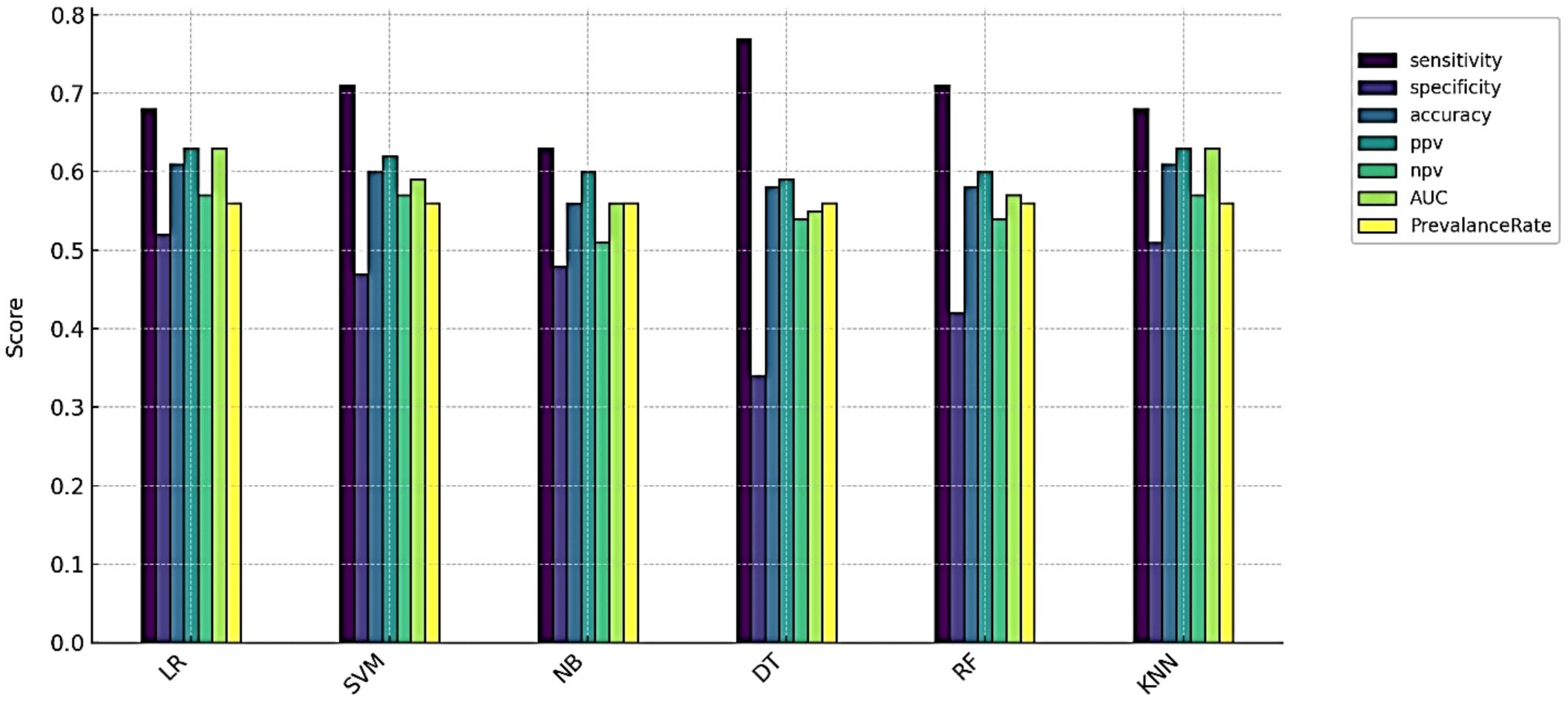
The results of various models (LR, SVM, NB, DT, and KNN) were assessed using sensitivity, specificity, PPV, NPV, AUC, and prevalence rate.

### Important Features

To assess the contribution of each descriptor type, we systematically removed one feature type at a time, and recalculated the evaluation metrics of the proposed prediction model. A greater decline in metrics indicated a higher importance of the removed feature type. As shown in figure 2, GC content, Gene Ontology semantic similarity, Normalized frequency of beta-structure, normalized frequency of alpha-helix, and coil emerge as the top four features, playing a critical role in predictive accuracy.

**Figure 2.**
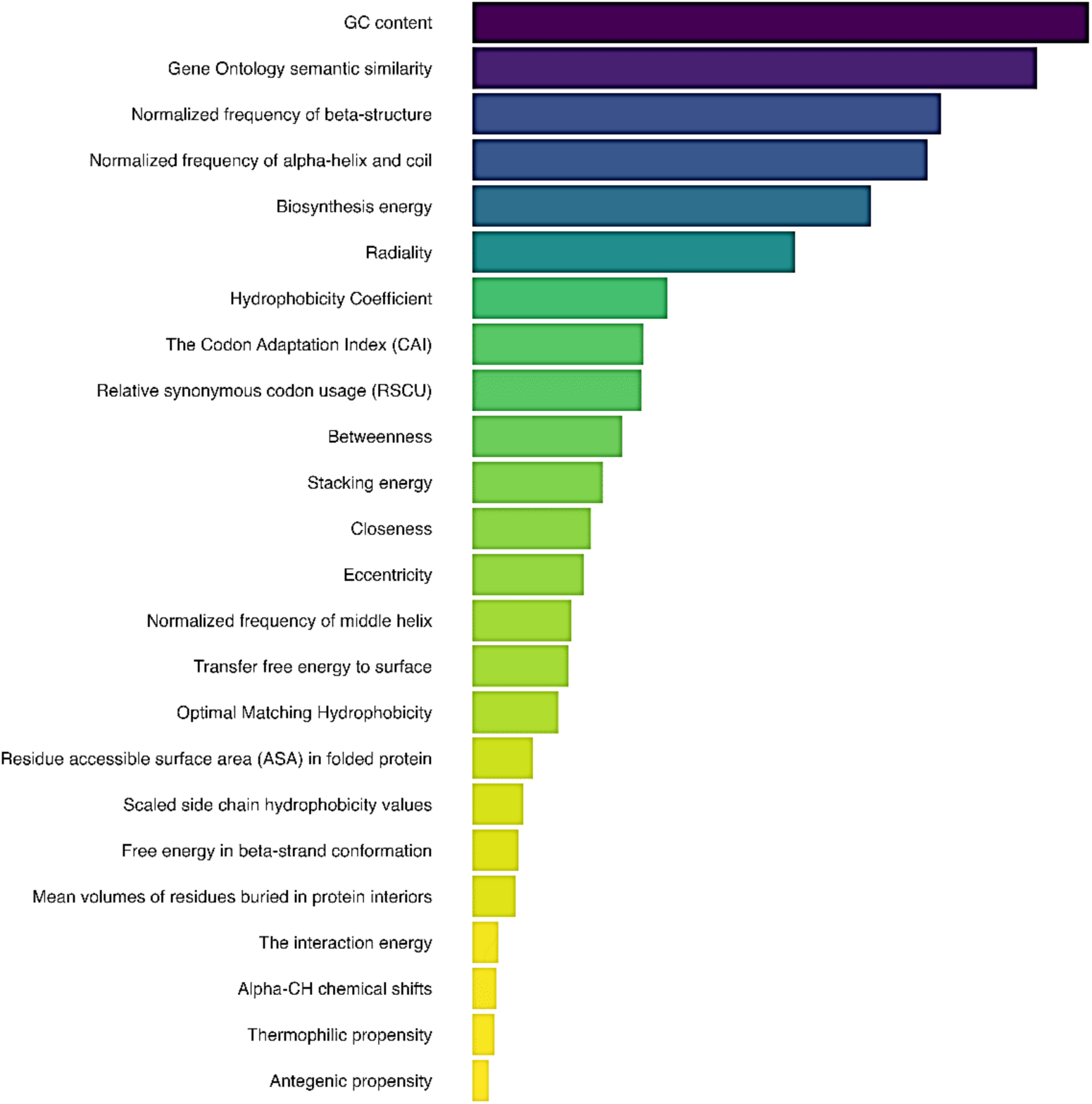
All features were ranked in order of importance, from the most significant to the least significant, and the top ten were selected for further analysis.

Features related to structural and biochemical properties, such as Biosynthesis energy, Radiality, and Hydrophobicity Coefficient, also rank high, underlining their biological relevance. Lower-ranked features, such as Antigenic propensity and Thermophilic propensity, contribute less significantly to model performance, validating their exclusion during dimensionality reduction.

Following the feature importance analysis, we trained the models using the selected key features and assessed their performance metrics. The radar chart (Figure 3.) illustrates a comparison between models trained on key feature sets and those trained on the full feature sets. The findings reveal that focusing on important features resulted in more accurate predictions for most classifiers, while maintaining efficiency comparable to models using all features. The performance remains consistent between all features and important features, with minimal deviation across all classifiers. This indicates that the selected important features capture the key patterns in the data effectively, offering equivalent predictive power as the full feature set.

**Figure 3.**
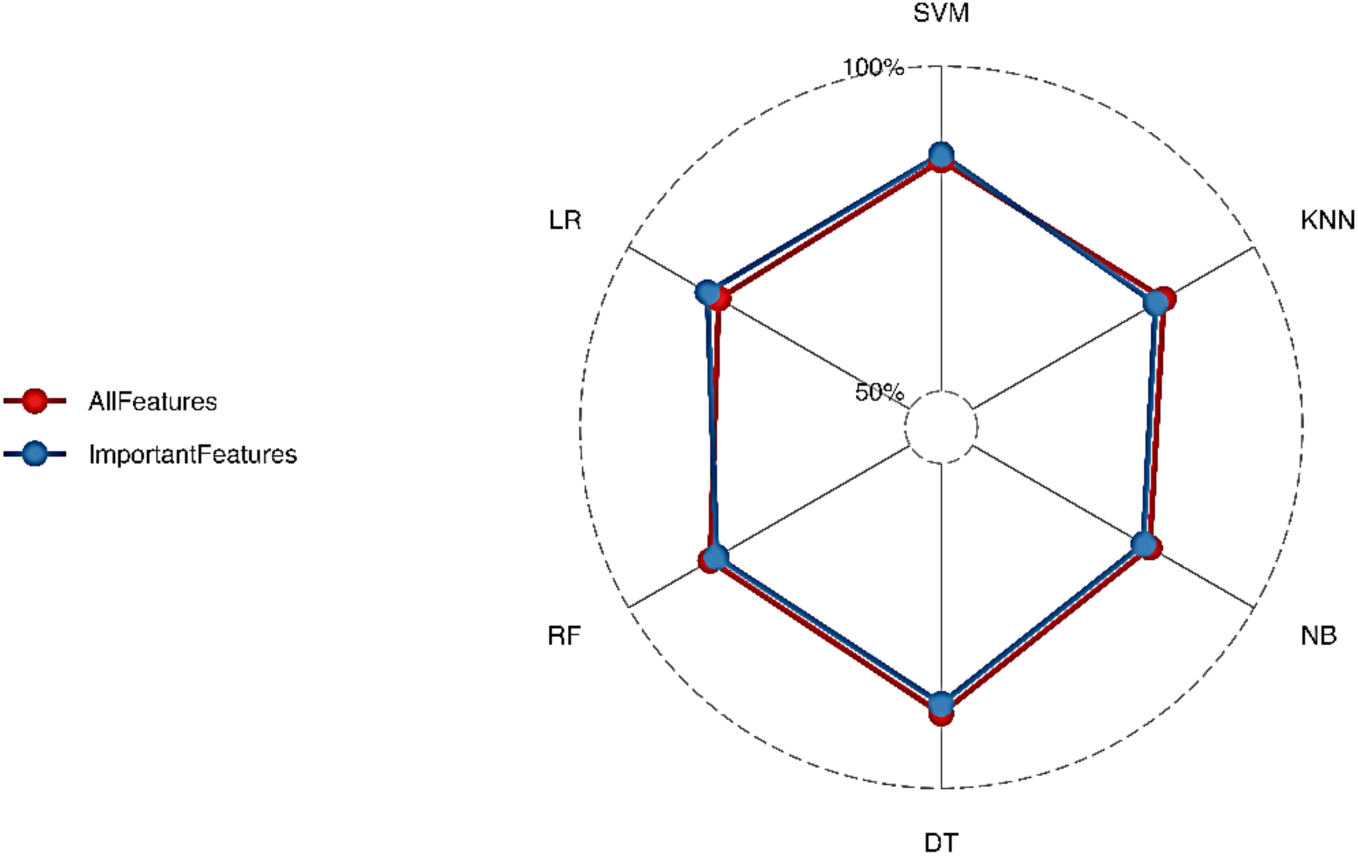
Radar graph to show the comparative performance of LR, SVM, KNN, NB, RF, and DT classifiers when trained on important features versus all features.

### Disease Enrichment Analysis

The disease enrichment analysis using the DisGeNET database in Enrichr identified significant associations between the virus-interacting host proteins and various cancer types, particularly liver carcinoma (p = 2.61 × 10⁻²⁷), which exhibited the highest statistical significance. Additional highly enriched disease terms included neoplasm metastasis (p = 7.32 × 10⁻²⁷) and tumor progression (p = 1.79 × 10⁻²⁰), suggesting that the identified proteins may be involved in cancer-related mechanisms, including tumor development and metastatic spread.

Moreover, Hepatitis C (p = 1.99 × 10⁻²²) and Hepatitis B (p = 2.91 × 10⁻¹⁹) were also significantly enriched, reinforcing the potential role of viral infections in liver-related pathologies. Since chronic infections with these viruses are well-documented risk factors for hepatocellular carcinoma, this enrichment suggests a functional overlap between the virus-interacting proteins in this study and those implicated in viral hepatitis-associated liver disease progression.

**Figure 4.**
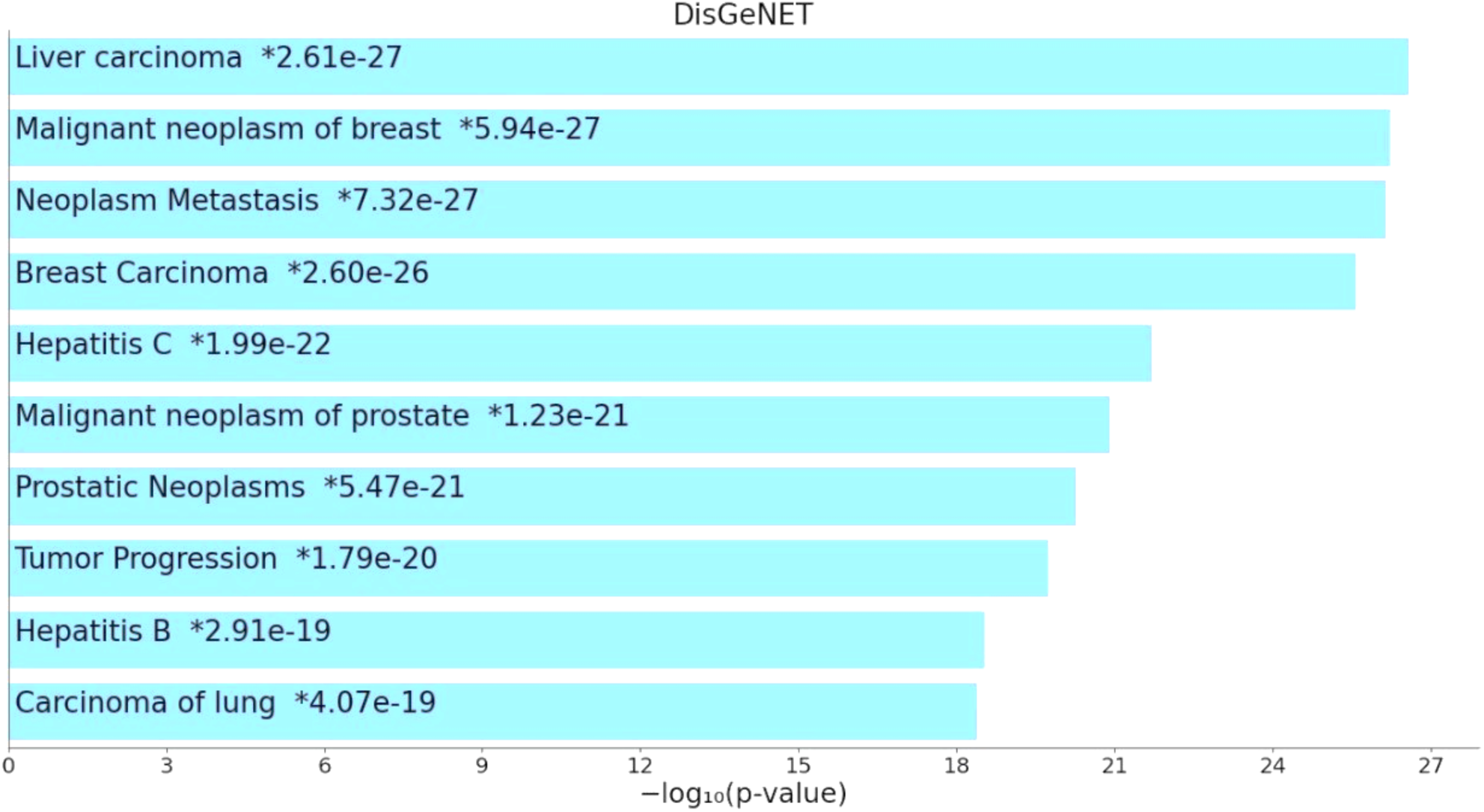
Bar chart of top enriched terms from the DisGeNET gene set library. The top enriched terms for the input gene set are displayed based on the lLog10(p-value), with the actual p-value shown next to each term. The term at the top has the most significant overlap with the input query gene set.

Interestingly, besides liver-related diseases, malignant neoplasms of breast (78) (p = 5.94 × 10⁻²⁷) and malignant neoplasms of the prostate (p = 1.23 × 10⁻²¹) were among the most significantly enriched disease categories (79). This may indicate that some of the identified proteins are also involved in broader oncogenic processes that extend beyond liver cancer, potentially playing a role in cell cycle regulation, apoptosis, or immune evasion mechanisms shared across multiple cancer types. Overall, these findings suggest that the virus-interacting host proteins identified in this study are not only functionally enriched in liver cancer pathways but also exhibit significant associations with metastatic progression and viral hepatitis, which are key factors in hepatocarcinogenesis. This highlights potential avenues for further investigation into the role of viral infections in cancer development and progression. Table 1 presents the genes enriched in liver cancer along with their functions.

**Table 1.**
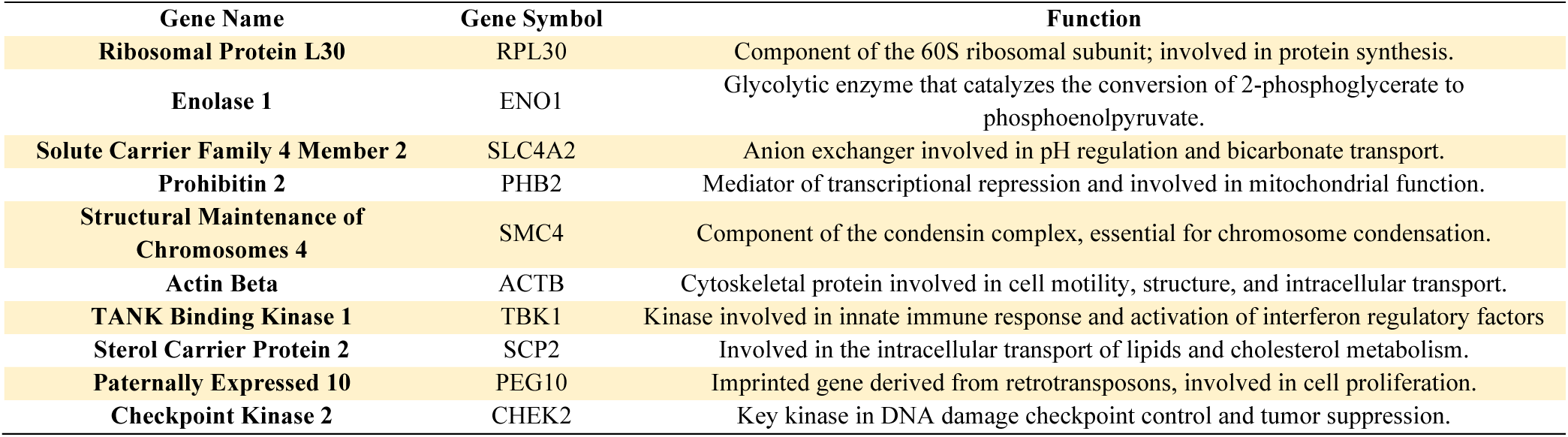
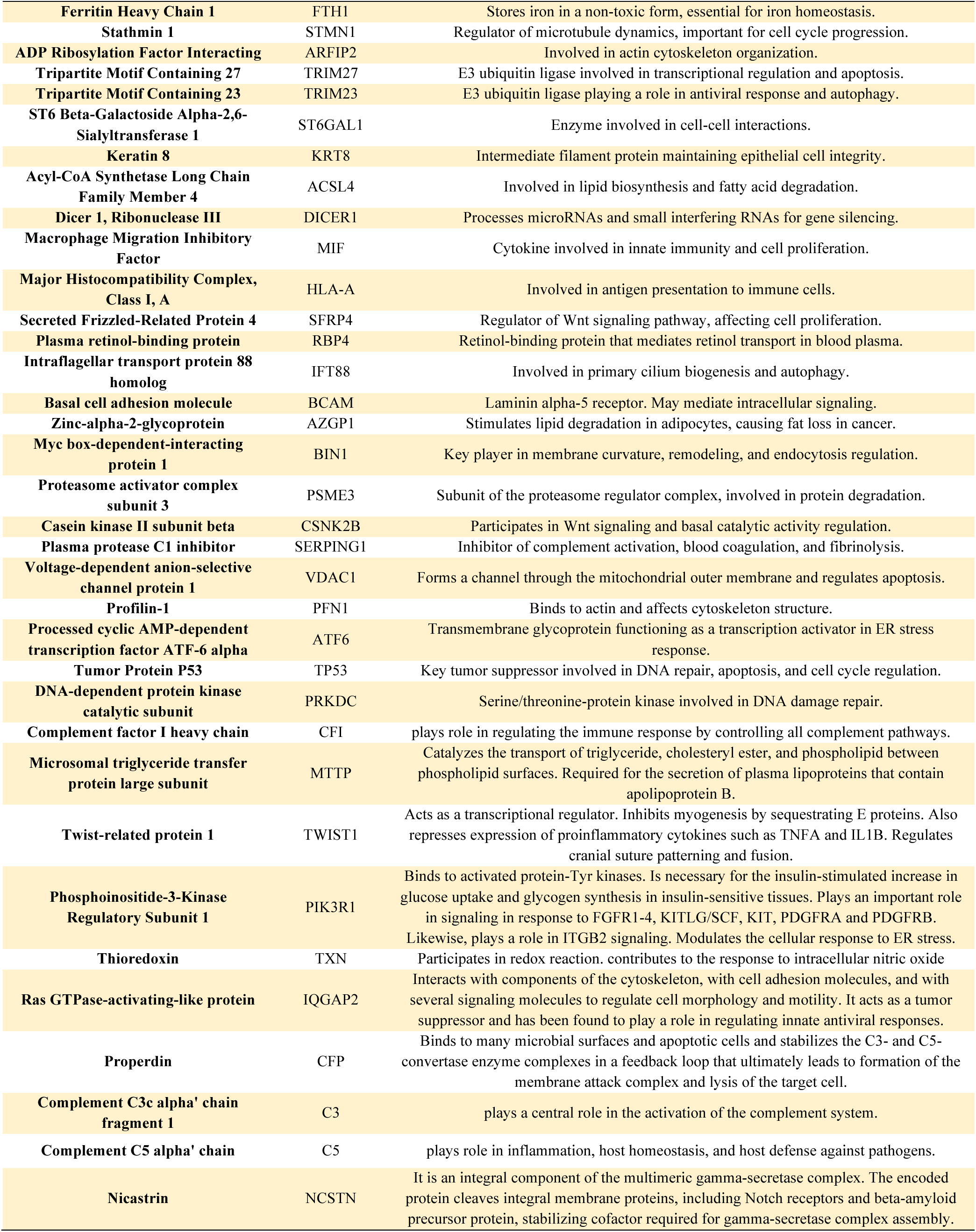
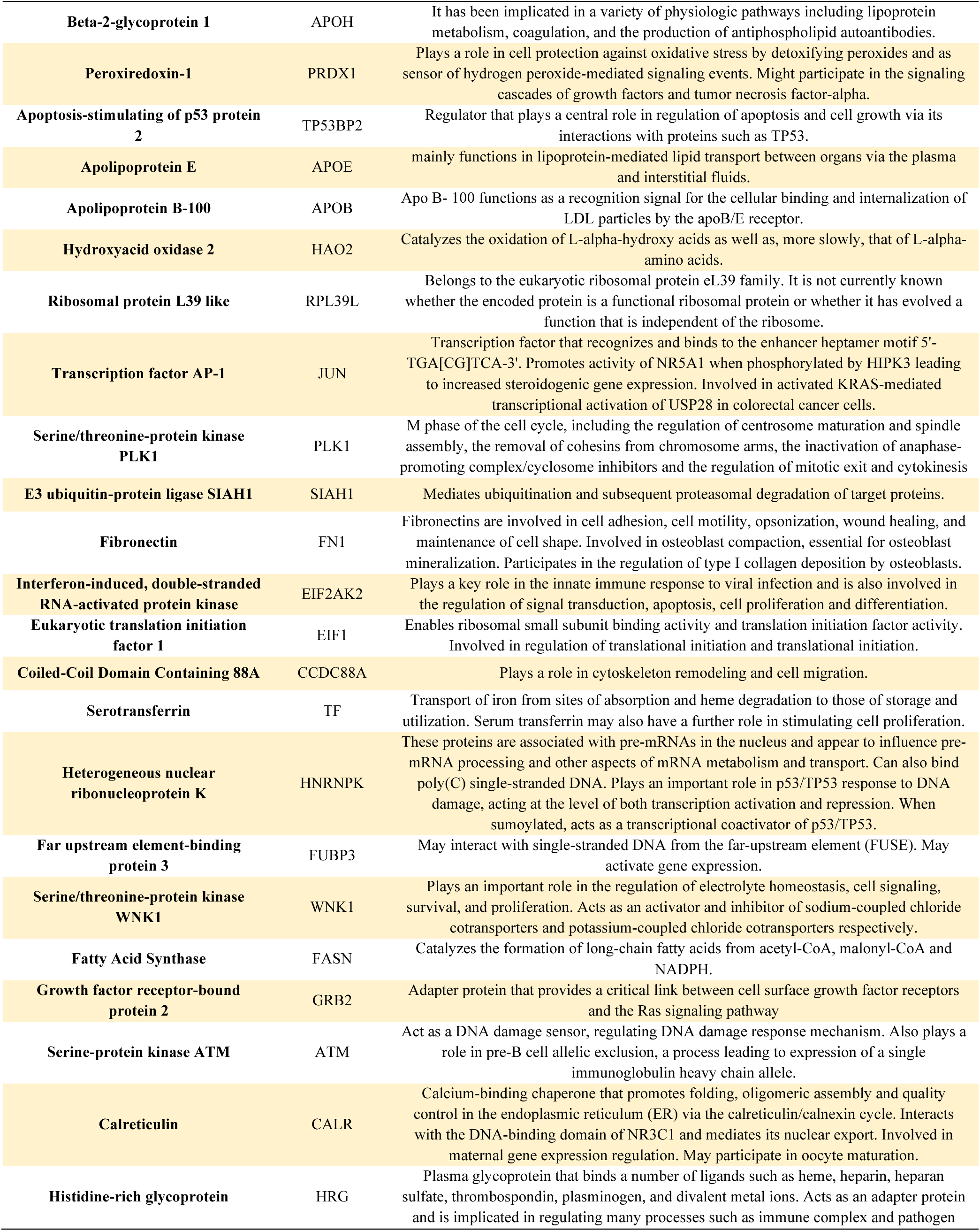
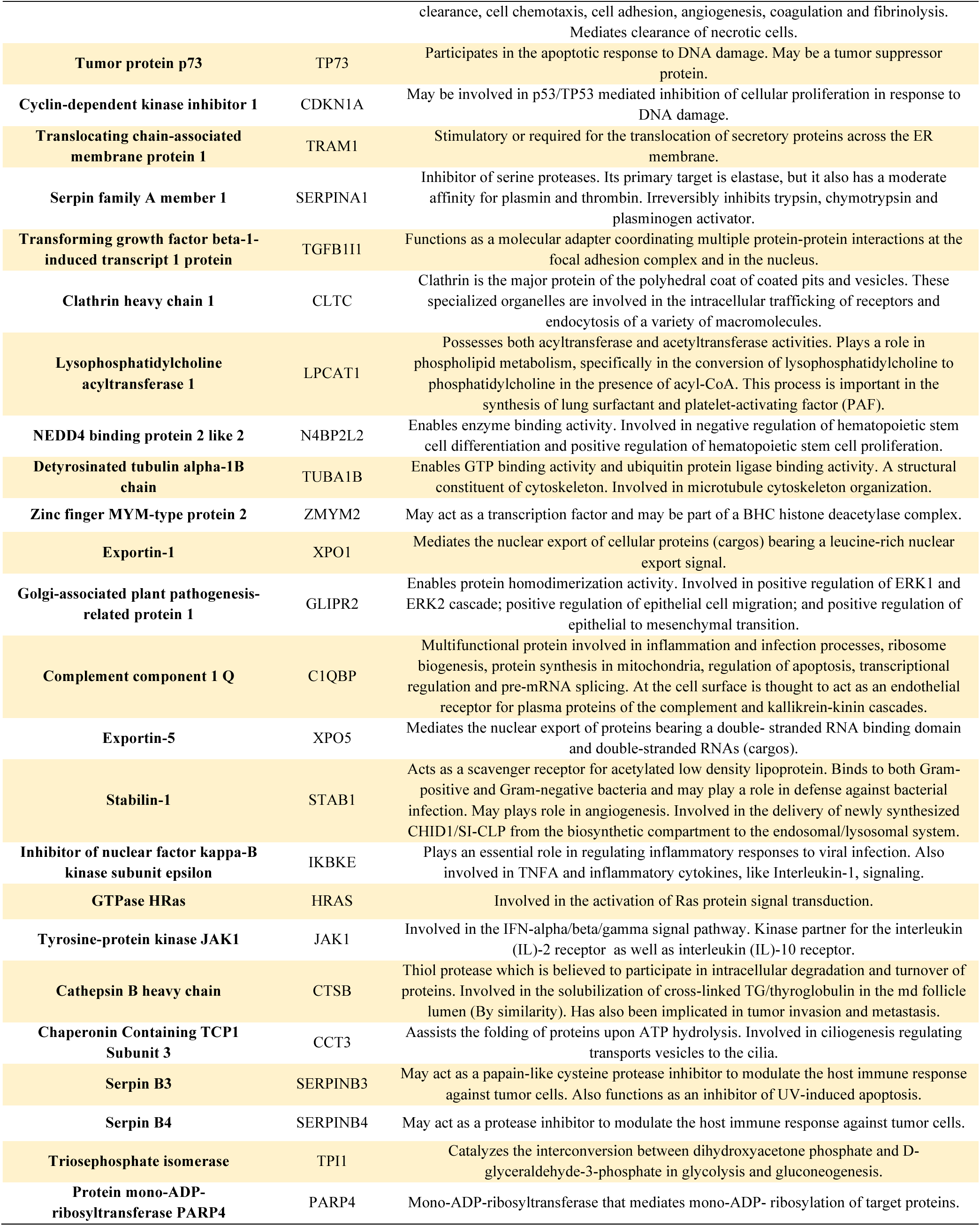
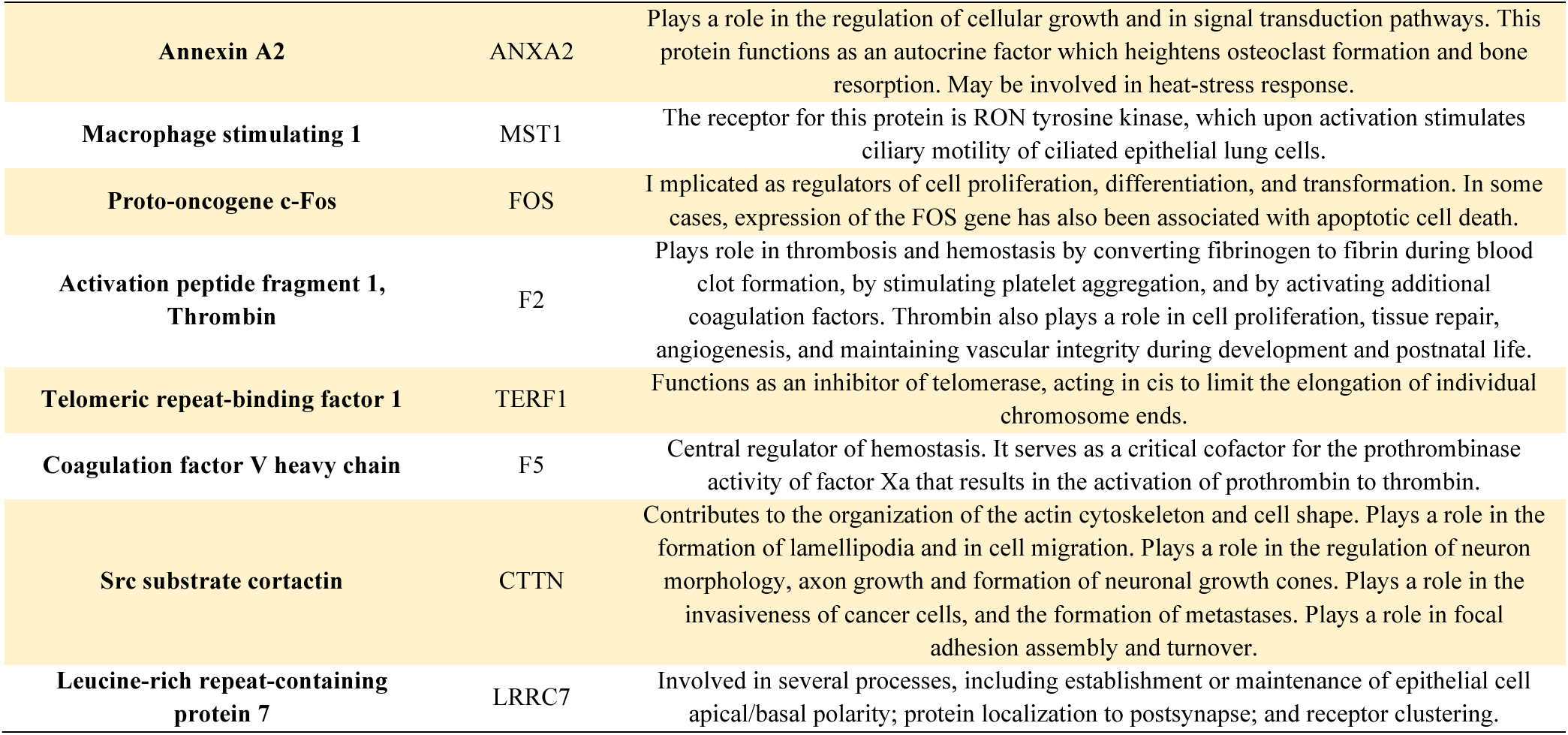
Hepatocellular Carcinoma-Associated Genes Identified Through Enrichment Analysis and Their Functions.

### GO Term Enrichment Analysis

To further investigate the molecular mechanisms underlying this association, we integrated functional enrichment analysis from Gene Ontology (GO) terms, using the DAVID (the database for database for annotation, visualization, and integrated discovery), with disease enrichment results. A total of 951 enriched molecular functions, 952 enriched cellular components, and 935 enriched biological processes were identified. The enrichment analysis highlights key cellular vulnerabilities exploited by the virus, revealing disruptions in chromatin organization, protein stability, and immune signaling. The most significant biological processes suggest viral interference in CENP-A chromatin assembly and heterochromatin organization, potentially altering host transcriptional control. Molecular function enrichment points to a strong reliance on host protein and RNA interactions, indicative of viral strategies to hijack cellular machinery for replication. At the cellular level, the virus appears to exploit exosome-mediated communication, possibly facilitating immune evasion and systemic spread. Pathway enrichment further reinforces a link to oncogenic processes, with signatures of endoplasmic reticulum stress, immune modulation, and metabolic shifts that may contribute to persistent infection and tumorigenesis. The GO enrichment analysis highlighted several functional categories that are biologically relevant to liver cancer development:

#### 1. Proteostasis and Endoplasmic Reticulum (ER) Stress

The enrichment of protein folding and protein stabilization, along with protein processing in the ER suggests that viral proteins may exploit host proteostasis pathways to evade degradation.

Cancer cells, particularly in HCC, rely on enhanced protein folding mechanisms to cope with metabolic stress, and viral manipulation of these pathways may promote tumor survival and immune evasion (80).

#### 2. Chromatin Organization and Telomere Stability

The enrichment of protein localization to CENP-A containing chromatin and heterochromatin organization suggests viral interference in epigenetic regulation, potentially altering host transcriptional programs. The enrichment of telomere organization further suggests that viral interactions may influence telomere stability, a key factor in cellular senescence and cancer progression. Telomere dysfunction is a known driver of HCC, as it contributes to genomic instability and uncontrolled proliferation, which are critical for tumor initiation (81).

#### 3. RNA Binding and Translational Control

The enrichment of RNA binding and protein binding suggests viral interference with host RNA metabolism, likely affecting gene expression and viral replication efficiency. In liver cancer, dysregulation of RNA-binding proteins (RBPs) is associated with oncogene activation, alternative splicing abnormalities, and immune evasion (82, 83).

#### 4. Inflammatory and Cell Death Pathways

The enrichment of neutrophil extracellular trap formation and necroptosis suggests a viral role in shaping the immune response and cell death mechanisms (84). Chronic inflammation and non-apoptotic cell death mechanisms such as necroptosis have been implicated in HCC progression, further supporting a connection between viral infection and pro-tumorigenic inflammation (85).

#### 5. Mitochondrial Quality Control and Mitophagy

The enrichment of mitophagy indicates viral modulation of mitochondrial turnover and metabolic adaptation. Mitophagy plays a dual role in liver cancer, preventing excessive oxidative stress in early tumorigenesis but later enabling tumor survival by removing damaged mitochondria. Viral regulation of mitophagy may facilitate metabolic reprogramming, allowing infected cells to adapt to stress while maintaining energy homeostasis (86). The significant enriched gene ontology terms and KEGG pathways are outlined in table 2.

**Table 2.** Enriched pathways and Gene Ontology (GO) terms identified in human proteins interacting with HEV, with significance determined by a Benjamini-corrected P-value threshold of less than 0.005.

| Category | Enriched feature | FDR |
| --- | --- | --- |
| Biological process | GO:0061644~protein localization to CENP-A containing chromatin | 3.32832254966858E-14 |
|  | GO:0006457~protein folding | 2.12722294878199E-10 |
|  | GO:0070828~heterochromatin organization | 4.17107969509387E-10 |
|  | GO:0032200~telomere organization | 5.82809530208538E-08 |
|  | GO:0070828~heterochromatin organization | 8.11398252391831E-08 |
|  | GO:0050821~protein stabilization | 1.25068263190799E-07 |
| Molecular function | GO:0005515~protein binding | 4.05998083208454E-47 |
|  | GO:0003723~RNA binding | 5.7682099654084E-39 |
|  | GO:0042802~identical protein binding | 4.060779610109531E-15 |
|  | GO:0046982~protein heterodimerization activity | 9.437589681332071E-15 |
|  | GO:0030527~structural constituent of chromatin | 5.27854694418556E-12 |
|  | GO:0051082~unfolded protein binding | 7.88111707287244E-12 |
| Cellular component | GO:0070062~extracellular exosome | 1.6343913724867799E-56 |
|  | GO:0005829~cytosol | 3.05738267468569E-33 |
|  | GO:0016020~membrane | 6.30745126373513E-20 |
|  | GO:0005788~endoplasmic reticulum lumen | 7.97372108526913E-19 |
|  | GO:0005654~nucleoplasm | 4.98558821612305E-17 |
|  | GO:0005634~nucleus | 1.10134180696361E-16 |
| Pathway | hsa04613:Neutrophil extracellular trap formation | 3.37130810300817E-12 |
|  | hsa04141:Protein processing in endoplasmic reticulum | 6.83092976782834E-11 |
|  | hsa04217:Necroptosis | 5.23420960298492E-08 |
|  | hsa05203:Viral carcinogenesis | 6.07809950705983E-08 |
|  | hsa04137:Mitophagy - animal | 0.0000460711373164893 |
|  | hsa03082:ATP-dependent chromatin remodeling | 0.0000655797631657286 |

## Discussion

Predicting protein interactions, especially in host-pathogen systems, is challenging, and even small improvements over random guessing are valuable. This is even harder for pathogens like HEV, which have not been studied as much as their global impact warrants, making it difficult to assess the accuracy of predictions.

HEV codes three or four open reading frames (ORFs) depending on the genotype. ORF1 encodes a multifunctional nonstructural polyprotein involved in viral replication, containing domains such as a methyltransferase (87), helicase, and RNA-dependent RNA polymerase. ORF2 codes for the capsid protein, essential for viral assembly and host immune interactions. ORF3 is a small phosphoprotein that regulates viral egress, immune modulation, and host signaling pathways. In genotype 1, an additional ORF4 enhances viral replication under stress conditions by interacting with the viral RNA polymerase.

Using affinity chromatography, immunoprecipitation, and two-hybrid assays, recent studies have revealed that HEV hijacks multiple human proteins to facilitate its replication and evade immune responses. HNRNPK and HNRNPA2B1 assist HEV polymerase and RNA processing, while SHARPIN and RNF5 modulate interferon signaling through ORF3. Additional host factors, such as FTL, TMEM154, FCGR2A, and STUB1, further highlight HEV’s interaction with cellular pathways like ubiquitination, viral entry, and immune modulation (88, 89).

As one of the few machine learning-based investigations in this area, our study builds on Barman et al. (14), who used SVM, Naïve Bayes, and Random Forest, achieving 74% accuracy and 83% specificity but lacked HEV-specific validation. Our expanded classifier set, including Decision Trees (DT) and Logistic Regression (LR), improved sensitivity (up to 77%) at the cost of specificity (max 52%), prioritizing novel interaction discovery over minimizing false positives.

Feature selection is another key distinction. Barman et al. relied on domain-domain associations and amino acid composition, whereas our model incorporates several descriptors, including GC content, gene ontology similarity, and structural attributes, enhancing biological relevance. This broader feature set likely contributes to higher sensitivity, making our approach more suited for exploratory HEV-host interaction discovery.

Furthermore, our evaluation framework extends beyond standard metrics, integrating prevalence rate, AUC, and interpretability analyses for a more biologically meaningful assessment. While Barman’s model is ideal for high-specificity predictions, ours excels in capturing novel interactions, emphasizing sensitivity for comprehensive PPI mapping.

To date, no study has systematically elucidated the molecular mechanisms by which HEV may contribute to hepatocarcinogenesis. The reviews by Klöhn et al. (2021) and Shen et al. (2023) highlight that although direct evidence for HEV-driven hepatocellular carcinoma is scarce, chronic HEV infection has been associated with liver fibrosis, cirrhosis, and immune dysregulation, all of which are recognized risk factors for HCC. Our study identified significant enrichment in pathways related to viral carcinogenesis, hepatitis B and C, and apoptosis, indicating that HEV-associated protein interactions may engage pathways known to contribute to liver disease progression. Furthermore, we identified a significant association between virus-interacting host proteins and liver carcinoma. Although epidemiological studies have not yet firmly established HEV as an independent oncogenic virus, the presence of overlapping host responses with established viral carcinogenesis pathways suggests that HEV infection could act as a cofactor in liver disease progression, particularly in patients with pre-existing liver conditions. The literature suggests that HEV may contribute to HCC by inhibiting the PI3K/AKT/mTOR pathway while modulating apoptosis and angiogenesis regulators. This includes Bax, Bcl-2, Apaf-1, caspase-3, caspase-9, and angiogenesis-related proteins DHX9 and TGFB, all of which are implicated in tumor progression (90).

Given the growing recognition of chronic HEV infections in immunocompromised individuals, further research is needed to determine whether persistent HEV infection alone contributes to hepatocarcinogenesis or primarily exacerbates underlying liver pathology. Its potential role in HCC progression raises concerns about the need for preventive measures in HEV-infected individuals.

## Conclusion

A comprehensive gold standard for human-HEV protein interactions is currently lacking. However, despite the limited known HEV-human interactions, our method demonstrates robust predictive capability, aligning with findings from existing studies. By broadening classifier diversity and enhancing biological interpretability, our study provides a more comprehensive framework for HEV-host protein-protein interaction (PPI) prediction. The predicted interactions shed light on potential viral mechanisms and highlight key pathways that warrant further investigation. Future experimental studies should focus on validating these host-virus interactions to determine their direct role in liver carcinogenesis and assess their therapeutic potential in virus-associated liver cancer.

## References

1. Peron J-M, Larrue H, Izopet J, Buti M. The pressing need for a global HEV vaccine. Journal of Hepatology. 2023.

2. Wang B, Meng X-J. Structural and molecular biology of hepatitis E virus. Computational and structural biotechnology journal. 2021;19:1907–16.

3. Adlhoch C, Avellon A, Baylis SA, Ciccaglione AR, Couturier E, de Sousa R, et al. Hepatitis E virus: Assessment of the epidemiological situation in humans in Europe, 2014/15. Journal of Clinical Virology. 2016;82:9–16.

4. De Schryver A, De Schrijver K, François G, Hambach R, van Sprundel M, Tabibi R, et al. Hepatitis E virus infection: an emerging occupational risk? Occupational medicine. 2015;65(8):667–72.

5. Lu L, Li C, Hagedorn CH. Phylogenetic analysis of global hepatitis E virus sequences: genetic diversity, subtypes and zoonosis. Reviews in medical virology. 2006;16(1):5–36.

6. Al-Shimari FH, Rencken CA, Kirkwood CD, Kumar R, Vannice KS, Stewart BT. Systematic review of global hepatitis E outbreaks to inform response and coordination initiatives. BMC Public Health. 2023;23(1):1120.

7. Klöhn M, Schrader JA, Brüggemann Y, Todt D, Steinmann E. Beyond the usual suspects: Hepatitis E virus and its implications in hepatocellular carcinoma. Cancers. 2021;13(22):5867.

8. Webb G, Dalton H. Hepatitis E: an expanding epidemic with a range of complications. Clinical Microbiology and Infection. 2020;26(7):828–32.

9. Purdy MA, Drexler JF, Meng X-J, Norder H, Okamoto H, Van der Poel WH, et al. ICTV virus taxonomy profile: Hepeviridae 2022. Journal of General Virology. 2022;103(9):001778.

10. Liu X, Lu Y, Wang L, Geng W, Shi X, Zhang X. RF-PSSM: A Combination of Rotation Forest Algorithm and Position-Specific Scoring Matrix for Improved Prediction of Protein-Protein Interactions Between Hepatitis C Virus and Human. Big Data Mining and Analytics. 2022;6(1):1–11.

11. Houri H, Aghdaei HA, Firuzabadi S, Khorsand B, Soltanpoor F, Rafieepoor M, et al. High prevalence rate of microbial contamination in patient-ready gastrointestinal endoscopes in Tehran, Iran: an alarming sign for the occurrence of severe outbreaks. Microbiology Spectrum. 2022;10(5):e01897–22.

12. Songtanin B, Molehin AJ, Brittan K, Manatsathit W, Nugent K. Hepatitis E Virus Infections: Epidemiology, Genetic Diversity, and Clinical Considerations. Viruses. 2023;15(6):1389.

13. Samandari Bahraseman MR, Khorsand B, Esmaeilzadeh-Salestani K, Sarhadi S, Hatami N, Khaleghdoust B, et al. The use of integrated text mining and protein-protein interaction approach to evaluate the effects of combined chemotherapeutic and chemopreventive agents in cancer therapy. Plos one. 2022;17(11):e0276458.

14. Barman RK, Saha S, Das S. Prediction of interactions between viral and host proteins using supervised machine learning methods. PloS one. 2014;9(11):e112034.

15. Evans P, Dampier W, Ungar L, Tozeren A. Prediction of HIV-1 virus-host protein interactions using virus and host sequence motifs. BMC medical genomics. 2009;2:1–13.

16. Ibrahim AH, Karabulut OC, Karpuzcu BA, Türk E, Süzek BE. A correlation coefficient-based feature selection approach for virus-host protein-protein interaction prediction. Plos one. 2023;18(5):e0285168.

17. Mohammad SM, Jetty D. Machine Learning Techniques for Sequence-Based Prediction of Viral-Host Interactions. International Journal of Emerging Technologies and Innovative Research (www jetir org| UGC and issn Approved), ISSN. 2021:2349–5162.

18. Planas-Iglesias J, Marin-Lopez MA, Bonet J, Garcia-Garcia J, Oliva B. iLoops: a protein– protein interaction prediction server based on structural features. Bioinformatics. 2013;29(18):2360–2.

19. Zheng N, Wang K, Zhan W, Deng L. Targeting virus-host protein interactions: Feature extraction and machine learning approaches. Current drug metabolism. 2019;20(3):177–84.

20. Khorsand B, Khammari A, Shirvanizadeh N, Zahiri J, Arab SS. OligoCOOL: a mobile application for nucleotide sequence analysis. Biochemistry and Molecular Biology Education. 2019;47(2):201–6.

21. Khorsand B, Savadi A, Naghibzadeh M. SARS-CoV-2-human protein-protein interaction network. Informatics in medicine unlocked. 2020;20:100413.

22. Khorsand B, Savadi A, Zahiri J, Naghibzadeh M. Alpha influenza virus infiltration prediction using virus-human protein-protein interaction network. Mathematical Biosciences and Engineering. 2020;17(4):3109–29.

23. Chakraborty A, Mitra S, Bhattacharjee M, De D, Pal AJ. Determining human-coronavirus protein-protein interaction using machine intelligence. Medicine in Novel Technology and Devices. 2023;18:100228.

24. Kawashima S, Pokarowski P, Pokarowska M, Kolinski A, Katayama T, Kanehisa M. AAindex: amino acid index database, progress report 2008. Nucleic acids research. 2007;36(suppl_1):D202–D5.

25. Zareei S, Khorsand B, Dantism A, Zareei N, Asgharzadeh F, Zahraee SS, et al. PeptiHub: a curated repository of precisely annotated cancer-related peptides with advanced utilities for peptide exploration and discovery. Database. 2024;2024:baae092.

26. Fauchère JL, Charton M, Kier LB, Verloop A, Pliska V. Amino acid side chain parameters for correlation studies in biology and pharmacology. International journal of peptide and protein research. 1988;32(4):269–78.

27. Soltanyzadeh M, Khorsand B, Baneh AA, Houri H. Clarifying differences in gene expression profile of umbilical cord vein and bone marrow-derived mesenchymal stem cells; a comparative in silico study. Informatics in Medicine Unlocked. 2022;33:101072.

28. Grantham R. Amino acid difference formula to help explain protein evolution. science. 1974;185(4154):862–4.

29. Levitt M. A simplified representation of protein conformations for rapid simulation of protein folding. Journal of molecular biology. 1976;104(1):59–107.

30. Bundi A, Wüthrich K. 1H-NMR parameters of the common amino acid residues measured in aqueous solutions of the linear tetrapeptides H-Gly-Gly-X-L-Ala-OH. Biopolymers: Original Research on Biomolecules. 1979;18(2):285–97.

31. Harpaz Y, Gerstein M, Chothia C. Volume changes on protein folding. Structure. 1994;2(7):641–9.

32. Engelman D, Steitz T, Goldman A. Identifying nonpolar transbilayer helices in amino acid sequences of membrane proteins. Annual review of biophysics and biophysical chemistry. 1986;15(1):321–53.

33. Muñoz V, Serrano L. Intrinsic secondary structure propensities of the amino acids, using statistical ϕ–ψ matrices: comparison with experimental scales. Proteins: Structure, Function, and Bioinformatics. 1994;20(4):301–11.

34. Black SD, Mould DR. Development of hydrophobicity parameters to analyze proteins which bear post-or cotranslational modifications. Analytical biochemistry. 1991;193(1):72–82.

35. Miyazawa S, Jernigan RL. Self-consistent estimation of inter-residue protein contact energies based on an equilibrium mixture approximation of residues. Proteins: Structure, Function, and Bioinformatics. 1999;34(1):49–68.

36. Wolfenden R, Andersson L, Cullis P, Southgate C. Affinities of amino acid side chains for solvent water. Biochemistry. 1981;20(4):849–55.

37. Rose GD, Geselowitz AR, Lesser GJ, Lee RH, Zehfus MH. Hydrophobicity of amino acid residues in globular proteins. Science. 1985;229(4716):834–8.

38. Karplus P, Schulz G. Prediction of chain flexibility in proteins: a tool for the selection of peptide antigens. Naturwissenschaften. 1985;72(4):212–3.

39. Vihinen M, Torkkila E, Riikonen P. Accuracy of protein flexibility predictions. Proteins: Structure, Function, and Bioinformatics. 1994;19(2):141–9.

40. Janin J, Wodak S, Levitt M, Maigret B. Conformation of amino acid side-chains in proteins. Journal of molecular biology. 1978;125(3):357–86.

41. Eisenberg D, McLachlan AD. Solvation energy in protein folding and binding. Nature. 1986;319(6050):199–203.

42. Chothia C. The nature of the accessible and buried surfaces in proteins. Journal of molecular biology. 1976;105(1):1–12.

43. Cornette JL, Cease KB, Margalit H, Spouge JL, Berzofsky JA, DeLisi C. Hydrophobicity scales and computational techniques for detecting amphipathic structures in proteins. Journal of molecular biology. 1987;195(3):659–85.

44. Sweet RM, Eisenberg D. Correlation of sequence hydrophobicities measures similarity in three-dimensional protein structure. Journal of molecular biology. 1983;171(4):479–88.

45. Radzicka A, Wolfenden R. Comparing the polarities of the amino acids: side-chain distribution coefficients between the vapor phase, cyclohexane, 1-octanol, and neutral aqueous solution. Biochemistry. 1988;27(5):1664–70.

46. Qian N, Sejnowski TJ. Predicting the secondary structure of globular proteins using neural network models. Journal of molecular biology. 1988;202(4):865–84.

47. Bull HB, Breese K. Surface tension of amino acid solutions: a hydrophobicity scale of the amino acid residues. Archives of biochemistry and biophysics. 1974;161(2):665–70.

48. Aurora R, Rosee GD. Helix capping. Protein Science. 1998;7(1):21–38.

49. Roseman MA. Hydrophilicity of polar amino acid side-chains is markedly reduced by flanking peptide bonds. Journal of molecular biology. 1988;200(3):513–22.

50. Wimley WC, White SH. Experimentally determined hydrophobicity scale for proteins at membrane interfaces. Nature structural biology. 1996;3(10):842–8.

51. Crawford JL, Lipscomb WN, Schellman CG. The reverse turn as a polypeptide conformation in globular proteins. Proceedings of the National Academy of Sciences. 1973;70(2):538–42.

52. Finkelstein A, Badretdinov AY, Ptitsyn O. Physical reasons for secondary structure stability: α-Helices in short peptides. Proteins: Structure, Function, and Bioinformatics. 1991;10(4):287–99.

53. O’Neil KT, DeGrado WF. A thermodynamic scale for the helix-forming tendencies of the commonly occurring amino acids. Science. 1990;250(4981):646–51.

54. Kim CA, Berg JM. Thermodynamic β-sheet propensities measured using a zinc-finger host peptide. Nature. 1993;362(6417):267–70.

55. Blaber M, Zhang X-j, Matthews BW. Structural basis of amino acid α helix propensity. Science. 1993;260(5114):1637–40.

56. Wilce MC, Aguilar M-I, Hearn MT. Physicochemical basis of amino acid hydrophobicity scales: evaluation of four new scales of amino acid hydrophobicity coefficients derived from RP-HPLC of peptides. Analytical chemistry. 1995;67(7):1210–9.

57. George RA, Heringa J. An analysis of protein domain linkers: their classification and role in protein folding. Protein engineering. 2002;15(11):871–9.

58. Sneath P. Relations between chemical structure and biological activity in peptides. Journal of theoretical biology. 1966;12(2):157–95.

59. Nagano K. Logical analysis of the mechanism of protein folding: I. Predictions of helices, loops and β-structures from primary structure. Journal of molecular biology. 1973;75(2):401–20.

60. Robson B, Suzuki E. Conformational properties of amino acid residues in globular proteins. Journal of molecular biology. 1976;107(3):327–56.

61. Lifson S, Sander C. Antiparallel and parallel β-strands differ in amino acid residue preferences. Nature. 1979;282(5734):109-11.

62. Fodje M, Al-Karadaghi S. Occurrence, conformational features and amino acid propensities for the π-helix. Protein engineering. 2002;15(5):353–8.

63. Nakashima H, Nishikawa K. The amino acid composition is different between the cytoplasmic and extracellular sides in membrane proteins. FEBS letters. 1992;303(2-3):141–6.

64. Kumar S, Tsai C-J, Nussinov R. Factors enhancing protein thermostability. Protein engineering. 2000;13(3):179–91.

65. Fukuchi S, Nishikawa K. Protein surface amino acid compositions distinctively differ between thermophilic and mesophilic bacteria. Journal of molecular biology. 2001;309(4):835–43.

66. Deng L, Nie W, Zhao J, Zhang J. A hybrid deep learning framework for predicting the protein-protein interaction between virus and host. 2021.

67. Charoenkwan P, Chotpatiwetchkul W, Lee VS, Nantasenamat C, Shoombuatong W. A novel sequence-based predictor for identifying and characterizing thermophilic proteins using estimated propensity scores of dipeptides. Scientific Reports. 2021;11(1):23782.

68. Kolaskar AS, Tongaonkar PC. A semi-empirical method for prediction of antigenic determinants on protein antigens. FEBS letters. 1990;276(1-2):172–4.

69. Acharya D, Dutta TK. Elucidating the network features and evolutionary attributes of intra-and interspecific protein–protein interactions between human and pathogenic bacteria. Scientific Reports. 2021;11(1):190.

70. Khorsand B, Savadi A, Naghibzadeh M. Parallelizing assignment problem with DNA strands. Iranian Journal of Biotechnology. 2020;18(1):e2547.

71. Khorsand B, Asadzadeh Aghdaei H, Nazemalhosseini-Mojarad E, Nadalian B, Nadalian B, Houri H. Overrepresentation of Enterobacteriaceae and Escherichia coli is the major gut microbiome signature in Crohn’s disease and ulcerative colitis; a comprehensive metagenomic analysis of IBDMDB datasets. Frontiers in cellular and infection microbiology. 2022;12:1015890.

72. Khorsand B, Savadi A, Naghibzadeh M. Comprehensive host-pathogen protein-protein interaction network analysis. BMC bioinformatics. 2020;21:1–22.

73. Hourfar H, Taklifi P, Razavi M, Khorsand B. Machine Learning-Driven Identification of Molecular Subgroups in Medulloblastoma via Gene Expression Profiling. Clinical Oncology. 2025:103789.

74. Khorsand B, Vaghf A, Salimi V, Zand M, Ghoreishi SA. Enhancing Ischemic Stroke Management: Leveraging Machine Learning Models for Predicting Patient Recovery After Alteplase Treatment. medRxiv. 2024:2024.11. 05.24316803.

75. Khorsand B, Rajabnia M, Jahanian A, Fathy M, Taghvaei S, Houri H. Enhancing the accuracy and effectiveness of diagnosis of spontaneous bacterial peritonitis in cirrhotic patients: a machine learning approach utilizing clinical and laboratory data. Advances in Medical Sciences. 2025;70(1):1–7.

76. Irankhah L, Khorsand B, Naghibzadeh M, Savadi A. Analyzing the performance of short-read classification tools on metagenomic samples toward proper diagnosis of diseases. Journal of bioinformatics and computational biology. 2024;22(5):2450012.

77. Haghzad T, Khorsand B, Razavi SA, Hedayati M. A computational approach to assessing the prognostic implications of BRAF and RAS mutations in patients with papillary thyroid carcinoma. Endocrine. 2024;86(2):707–22.

78. Shiralipour A, Khorsand B, Jafari L, Salehi M, Kazemi M, Zahiri J, et al. Identifying key lysosome-related genes associated with drug-resistant breast cancer using computational and systems biology approach. Iranian Journal of Pharmaceutical Research: IJPR. 2022;21(1):e130342.

79. Razavi SA, Khorsand B, Salehipour P, Hedayati M. Metabolite signature of human malignant thyroid tissue: A systematic review and meta-analysis. Cancer Medicine. 2024;13(8):e7184.

80. Pavlović N, Heindryckx F. Exploring the role of endoplasmic reticulum stress in hepatocellular carcinoma through mining of the human protein atlas. Biology. 2021;10(7):640.

81. Ningarhari M, Caruso S, Hirsch TZ, Bayard Q, Franconi A, Védie A-L, et al. Telomere length is key to hepatocellular carcinoma diversity and telomerase addiction is an actionable therapeutic target. Journal of Hepatology. 2021;74(5):1155–66.

82. Zhang K, Barry AE, Lamm R, Patel K, Schafer M, Dang H. The role of RNA binding proteins in hepatocellular carcinoma. Advanced Drug Delivery Reviews. 2022;182:114114.

83. Zhou C, Wu Q, Zhao H, Xie R, He X, Gu H. Unraveling the role of RNA-binding proteins, with a focus on RPS5, in the malignant progression of hepatocellular carcinoma. International Journal of Molecular Sciences. 2024;25(2):773.

84. Kharaghani AA, Harzandi N, Khorsand B, Rajabnia M, Kharaghani AA, Houri H. High prevalence of Mucosa-Associated extended-spectrum β-Lactamase-producing Escherichia coli and Klebsiella pneumoniae among Iranain patients with inflammatory bowel disease (IBD). Annals of Clinical Microbiology and Antimicrobials. 2023;22(1):86.

85. Li X, Dong G, Xiong H, Diao H. A narrative review of the role of necroptosis in liver disease: a double-edged sword. Annals of Translational Medicine. 2021;9(5):422.

86. Ma X, McKeen T, Zhang J, Ding W. Role and mechanisms of mitophagy in liver diseases. Cells. 2020. Internet]; 2020.

87. Sadeghnezhad E, Sharifi M, Zare-maivan H, Khorsand B, Zahiri J. Cross talk between energy cost and expression of Methyl Jasmonate-regulated genes: from DNA to protein. Journal of Plant Biochemistry and Biotechnology. 2019;28:230–43.

88. Corneillie L, Lemmens I, Weening K, De Meyer A, Van Houtte F, Tavernier J, et al. Virus–Host Protein Interaction Network of the Hepatitis E Virus ORF2-4 by Mammalian Two-Hybrid Assays. Viruses. 2023;15(12):2412.

89. Kanade GD, Pingale KD, Karpe YA. Protein interactions network of hepatitis E virus RNA and polymerase with host proteins. Frontiers in microbiology. 2019;10:2501.

90. Yin X, Kan F. Hepatitis E virus infection and risk of hepatocellular carcinoma: A systematic review and meta-analysis. Cancer Epidemiology. 2023;87:102457.

